# Unsupervised deep learning on biomedical data with BoltzmannMachines.jl

**DOI:** 10.1101/578252

**Authors:** Stefan Lenz, Moritz Hess, Harald Binder

## Abstract

Deep Boltzmann machines (DBMs) are models for unsupervised learning in the field of artificial intelligence, promising to be useful for dimensionality reduction and pattern detection in clinical and genomic data. Multimodal and partitioned DBMs alleviate the problem of small sample sizes and make it possible to combine different input data types in one DBM model. We present the package “BoltzmannMachines” for the Julia programming language, which makes this model class available for practical use in working with biomedical data.

**Availability:** Notebook with example data: http://github.com/stefan-m-lenz/BMs4BInf2019 Julia package: http://github.com/stefan-m-lenz/BoltzmannMachines.jl

## 1 Motivation

In the field of artificial intelligence, generative models can learn potentially complex structure in biomedical data in an unsupervised manner. Subsequently, new, synthetic samples can be generated that represent what kind of biological structure has been uncovered. We focus on Deep Boltzmann Machines (DBMs) (Srivastava and Salakhutdinov, 2014) because these allow for flexible conditional sampling, and we have already adapted them for training with small sample sizes (Hess et al., 2017). While our previous approach was primarily designed for handling genetic data, our Julia package “BoltzmannMachines” now can integrate data of different types, i.e. multimodal data, e. g. clinical, gene expression, and single nucleotide polymorphism (SNP) data. We also provide convenient tools for the notorious challenge of hyperparameter tuning in our instance of deep learning.

In contrast to generative adversarial networks (GANs) (Goodfellow et al., 2014) and variational autoencoders (VAEs) (Kingma and Welling, 2013; Rezende et al., 2014), which are other popular generative models, DBMs aim at modeling the probability distribution and then make it easy to sample conditionally and access hidden nodes. While VAEs and GANs are trained based on a backpropagation algorithm, and thus can readily be implemented in frameworks such as TensorFlow (Abadi et al., 2015), DBMs require a different approach (see below). So we decided to build a specific tool in the Julia programming language (Bezanson et al., 2017), which allows for gradually implementing and optimizing algorithms and DBM architectures without having to implement time-critical parts in a low-level language such as C.

## 2 Features

In the following, we describe the most important features of our package, which are also illustrated with an exemplary multimodal application and published as a Jupyter notebook. The features are described in greater detail in the package documentation.

DBMs can be used for dimensionality reduction in a similar way as common techniques such as principal component analysis (PCA) or t-distributed stochastic neighbor embedding (t-SNE). This is done by training a DBM with a small number of nodes in the highest abstraction layer and plotting their activations against each other (see figure 2, panel A).

**Figure 1.**
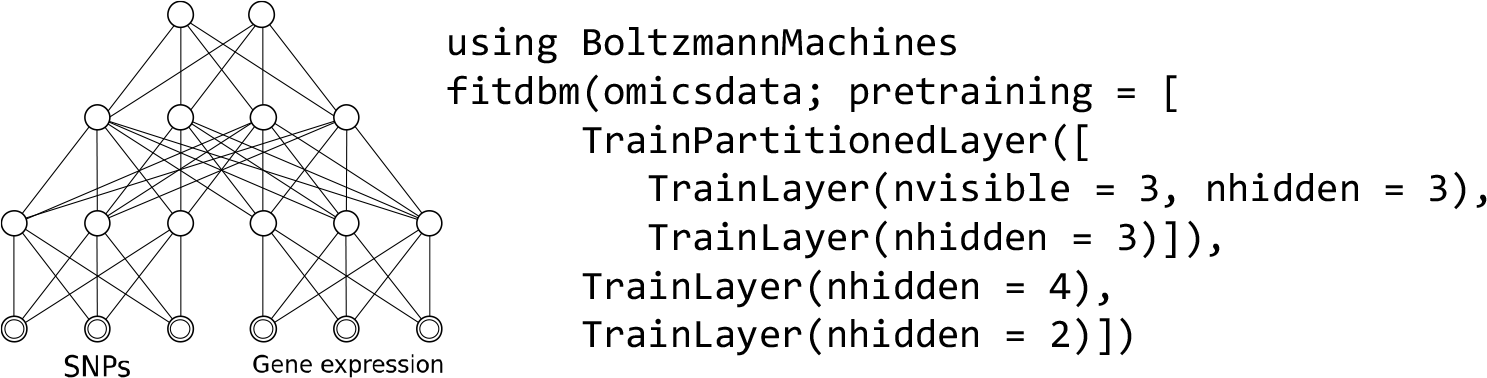
An exemplary architecture of a multimodal DBM and Julia code for creating the architecture.

**Figure 2.**
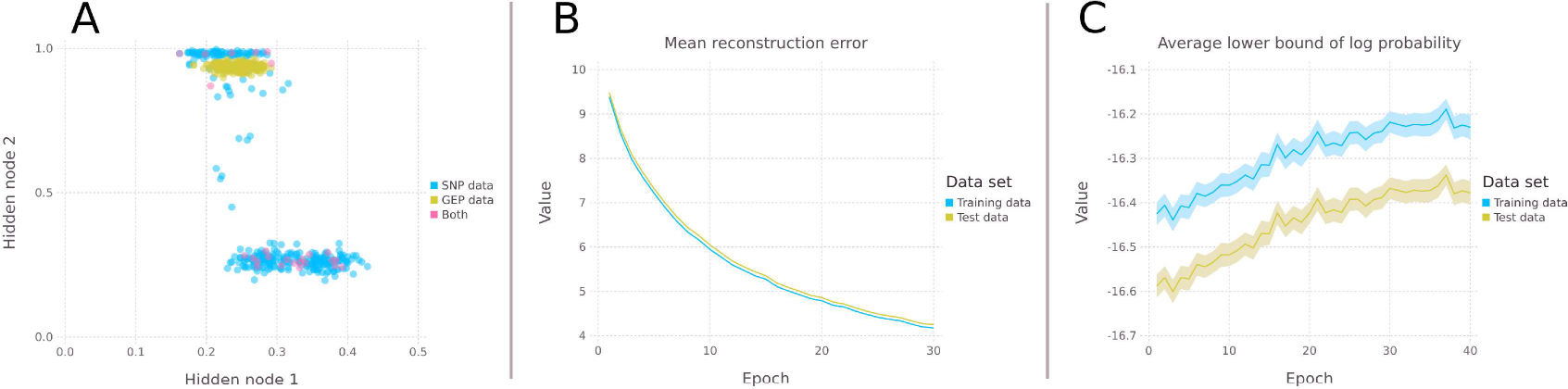
Plots created with the accompanying plotting package. Panel A shows a dimensionality reduction of multimodal data comprising of SNP and gene expression (GEP) data via the DBM’s hidden nodes. Panels B and C show exemplary monitoring ouput. The mean reconstruction error (panel B) should go down when pre-training an RBM layer, the estimated lower bound of the log-likelihood (panel C) should go up during DBM training until an optimum is reached. The uncertainty of the estimations is indicated by the ribbons around the graphs.

The training procedure implemented for DBMs consists of two parts: In the first step, a stack of restricted Boltzmann machines (RBMs) is trained. This then gives a good starting point for the subsequent training procedure employing mean-field approximation (Salakhutdinov and Hinton, 2009).

Due to low sample sizes, training deep learning models on genomic data poses a challenge. Compared to the number of measurements per patient or sample, e. g. SNPs or expressed genes, the number of available genome sequences or gene expression profiles is still low. But fewer samples can suffice for the training if the number of network parameters is reduced. In DBMs this can be done by grouping variables into blocks and by allowing connections between these blocks only at a higher level, thereby partitioning the network. Such parsimony in parameters is one use case for so-called multimodal DBMs, which are implemented in our package. Grouping variables may also be a way to incorporate background knowledge such as gene locations. It is natural in case of different measurements modalities such as SNP data and gene expression data or brain scans and test scores, which may also have different data types and/or may include measurement-specific patterns. To make the composition of such multimodal DBMs convenient and intuitive, the design of our package uses RBMs as simple building blocks for DBMs. It is possible to plug in different types of RBMs in the input layer, stack RBMs on top of each other to get a deeper network, and put RBMs side-by-side in partitioned layers. Different RBM types are available for modeling binary, continuous and categorical data. The package is also designed to enable fast development and integration of new types of RBMs at the input layer. The complete network architecture can be defined and trained in a single statement (see figure 1), which performs both layer-wise pre-training and the subsequent fine-tuning of the model. For pre-training, the hyperparameters of each RBM may be specified independently via TrainLayer arguments. It is also possible to individually monitor the training of RBM layers (see figure 2, panel B). An established optimization metric for this is the reconstruction error (Hinton, 2012), which is a proxy metric for the likelihood and computationally very easy to evaluate. After the stack of RBMs has been created, the composed network architecture can be trained and evaluated as a DBM.

The optimization target for DBM training is the variational lower bound of the model’s log-likelihood (see figure 2, panel C). The exact value for the likelihood can only be calculated exactly for very small models. For larger models, it can only be estimated by stochastic algorithms such as annealed importance sampling. Although the likelihood is hard to evaluate, it often remains as the only evaluation criterion for clinical or genomic data since other evaluation strategies such as judging generated images or sentences, which is popular for imaging data or in natural language processing, are not applicable. Exact likelihood calculation and estimation are both implemented in our package for all architectures of multimodal DBMs. This allows for monitoring the training process to see whether the optimization is successful. To additionally support other evaluation strategies, we designed the monitoring to be as easily extensible as possible. The training can thereby be examined from all angles. The optimization method itself is also designed to be customizable to run and evaluate different optimization algorithms.

